# ODELAM: Rapid sequence-independent detection of drug resistance in clinical isolates of Mycobacterium tuberculosis

**DOI:** 10.1101/2020.03.17.995480

**Authors:** Thurston Herricks, Magdalena Donczew, Fred D. Mast, Tige Rustad, Robert Morrison, Timothy R. Sterling, David R. Sherman, John D. Aitchison

## Abstract

Antimicrobial-resistant Mycobacterium tuberculosis (Mtb) causes over 200,000 deaths each year. Current assays of antimicrobial resistance need knowledge of mutations that confer drug resistance, or long periods of culture time to test growth under drug pressure. We present ODELAM (One-cell Doubling Evaluation of Living Arrays of Mycobacterium), a time-lapse microscopy-based method that observes individual cells growing into microcolonies. ODELAM enables rapid quantitative measures of growth kinetics in as little as 30 hours under a wide variety of environmental conditions. We show the utility of ODELAM by identifying ofloxacin resistance in clinical isolates of Mtb and benchmark its performance with standard MIC assays. In one clinical isolate, ODELAM identified ofloxacin heteroresistance and identifies the presence of drug resistant colony forming units (CFU) at 1 per 1000 CFUs in as little as 48 hours. ODELAM is a powerful new tool that can rapidly evaluate Mtb drug resistance in a laboratory setting.

## Introduction

The continued and accelerated emergence of anti-microbial resistance is a global threat and antimicrobial-resistant infections caused by *Mycobacterium tuberculosis* (Mtb) are particularly concerning. In 2018, approximately 484,000 people developed multidrug-resistant tuberculosis and an estimated 214,000 people died from rifampicin-resistant or multidrug-resistant tuberculosis (Geneva: World Health Organization 2019). Approximately 10% of all multidrug resistant tuberculosis infections are resistant to at least four commonly used anti-TB drugs (Dorman et al. 2018).

Rapid, reliable diagnosis is a key to the prevention and effective treatment of antimicrobial resistant infections (Boolchandani, D’Souza, and Dantas 2019). In the case of Mtb, culture-based methods remain the gold standard for identifying sensitivity to any antibiotic, but these methods require approximately two to four weeks (Kim 2005). Biomarker methods, such as the PCR-based Gene-Xpert assay, can rapidly identify antimicrobial resistant Mtb infections but require knowledge of specific genetic markers associated with resistance and are thus insensitive to unknown resistance mechanisms (Dorman et al. 2018). Detecting antimicrobial-resistant Mtb infections is especially challenging when bacterial sub-populations are present that possess differing levels of antimicrobial sensitivity (El-Halfawy and Valvano 2015). If present in relatively low abundance, these heteroresistant populations may evade detection until after treatment begins. Failure to rapidly identify and treat an antimicrobial-resistant or heteroresistant Mtb infection contributes to disease progression, treatment failure and tuberculosis relapse (Shin et al. 2018).

We developed ODELAM (One-cell Doubling Evaluation of Living Arrays of *Mycobacterium*), a time-lapse microscopy-based method designed to quantify growth phenotypes of populations of individual Mtb cells and colony forming units. Generally imaging platforms used to study mycobacteria use either microfluidic techniques or growth on solid media. Microfluidic platforms provide high resolution phenotypic analysis of mycobacterial cell biology but are limited in the number of cells that can be readily observed (Aldridge et al. 2012; Golchin et al. 2012; Wakamoto et al. 2013). Conventional assays on solid media provide crude growth analysis, including antibiotic sensitivity, on large populations of cells but require up to 8 weeks for observations whereas specialized microfluidic methods require on the order of 9 days (Choi et al. 2014).

ODELAM bridges these disparate methods by assaying growth temporally at a mesoscale on up to ∼100,000 CFUs in a single experiment. Originally designed for yeast, the adaptation offers improvements on phenotypic analysis and diagnostics for detecting and characterizing populations of drug resistant Mtb (Herricks et al. 2017). While commercial time-lapse and laboratory based genomic screening tools are available (Lee et al. 2019; Baryshnikova et al. 2010; Bean et al. 2014; Zackrisson et al. 2016; Golchin et al. 2012), ODELAM uniquely enables rapid quantitative measurements of growth kinetics of homogeneous and heterogeneous clinical isolates of Mtb in as little as 30 hrs and under a variety of environmental conditions, including antimicrobial drug pressures.

We demonstrate ODELAM by analyzing ofloxacin (OFX) resistance in clinical Mtb isolates. ODELAM was first benchmarked with the laboratory strain H37Rv and then two clinical isolates were tested, an OFX-resistant isolate and an isolate with OFX heteroresistance (Eilertson et al. 2016). Microscopy-based kinetic growth analysis of these strains under increasing drug pressures revealed characteristic growth and drug sensitivity phenotypes of each strain and specifically demonstrate the ability to rapidly detect heteroresistant sub-populations in a clinical isolate. ODELAM is a powerful new tool that can enable the timely identification of Mtb antimicrobial resistant phenotypes in a laboratory setting.

## Results

ODELAM is engineered to quantitatively assess growth kinetics of large populations of individual Mtb colony forming units (CFU). In each region of interest ODELAM can observe up to 1500 CFU and quantify their morphological and multiparameter growth phenotypes. A typical experiment involves 80 or 96 regions of interest. Single bacilli are observed over time as they grow into microscopically observable colonies (**Fig. 1A**). In these experiments, Mtb bacilli were observed to grow immediately after spotting on media. Mtb CFU were recorded for a total of 96 hrs and growth curves were extracted from the projected area of the individual colonies over time and fitted to the Gompertz function (**Fig. 1B**) (Herricks 2017). The kinetic parameters doubling time (Td), lag time (T-Lag), time in exponential phase (T-Exp) and number of doublings (Num Dbl) were derived from the Gompertz function and plotted as normalized frequency histograms which revealed population distributions (**Fig. 1C**). Summary non-parametric statistics for the isolates measured are presented as interquartile range (IQR) with respect to the median (**Table I**). For example, H37Rv, an Mtb strain commonly used in research laboratories, had a median lag time of 0.5 sec and grew with a doubling time median of 22.9 hrs (IQR 7.8). H37Rv bacilli grew in exponential phase for a median of 52.5 hrs (IQR 33.8) and during this time underwent a median of 3.1 doublings (IQR 1.6). These data are consistent with bulk laboratory measurements (Peñuelas-Urquides et al. 2013) and demonstrate the innate heterogeneity of the genetically homogeneous clonal population.

**Table I:** Summary growth statistics of isolates.

| Condition tested | CFU Total | Num of Rep | Med Td (Hrs) | IQR Td (Hrs) | Med T-Lag Time (Hrs) | IQR T-Lag Time (Hrs) | Med T-Exp (Hrs) | IQR T-Exp (Hrs) | Med Num Dbl | IQR Num Dbl |
| --- | --- | --- | --- | --- | --- | --- | --- | --- | --- | --- |
| Control-H37Rv | 5413 | 8 | 22.9 | 7.8 | 0.0 | 3.7 | 52.5 | 33.8 | 3.1 | 1.6 |
| 0.0625 ug/mL-H37Rv | 3380 | 4 | 23.4 | 9.6 | 0.0 | 2.8 | 51.8 | 28.1 | 3.0 | 1.5 |
| 0.125 ug/mL-H37Rv | 2946 | 4 | 24.3 | 10.0 | 0.0 | 2.9 | 53.0 | 29.5 | 3.0 | 1.4 |
| 0.5 ug/mL-H37Rv | 2231 | 4 | 22.6 | 10.7 | 0.9 | 6.2 | 35.2 | 16.2 | 2.0 | 0.9 |
| 2 ug/mL-H37Rv | 5354 | 8 | 19.2 | 13.7 | 3.9 | 6.1 | 17.3 | 8.2 | 0.9 | 0.6 |
| 4 ug/mL-H37Rv | 1412 | 4 | 21.1 | 12.5 | 3.9 | 5.2 | 14.7 | 7.7 | 0.6 | 0.4 |
| 8 ug/mL-H37Rv | 1728 | 4 | 20.8 | 12.1 | 2.8 | 4.0 | 11.5 | 6.5 | 0.5 | 0.3 |
| 16 ug/mL-H37Rv | 894 | 4 | 22.9 | 24.4 | 1.5 | 4.4 | 11.2 | 12.4 | 0.5 | 0.3 |
| Control-TRS10 | 3449 | 8 | 28.4 | 8.5 | 2.8 | 9.3 | 79.7 | 54.2 | 3.5 | 2.5 |
| 0.0625 ug/mL-TRS10 | 2181 | 4 | 28.2 | 6.9 | 1.9 | 8.1 | 81.7 | 55.6 | 3.8 | 2.6 |
| 0.125 ug/mL-TRS10 | 2014 | 4 | 28.5 | 7.6 | 0.0 | 5.4 | 80.3 | 61.2 | 3.7 | 2.8 |
| 0.5 ug/mL-TRS10 | 1167 | 4 | 28.3 | 6.3 | 0.0 | 6.0 | 81.8 | 62.5 | 3.8 | 3.1 |
| 2 ug/mL-TRS10 | 2982 | 8 | 30.6 | 17.9 | 2.3 | 9.2 | 73.7 | 42.6 | 2.8 | 2.2 |
| 4 ug/mL-TRS10 | 1190 | 4 | 53.1 | 26.0 | 0.0 | 7.4 | 65.7 | 38.7 | 1.5 | 1.1 |
| 8 ug/mL-TRS10 | 1063 | 4 | 57.8 | 28.4 | 0.0 | 2.1 | 56.7 | 27.4 | 1.3 | 0.7 |
| 16 ug/mL-TRS10 | 966 | 4 | 46.0 | 21.8 | 0.0 | 13.8 | 61.8 | 44.8 | 1.6 | 1.3 |
| Control-ADB42 | 5599 | 8 | 24.4 | 7.8 | 1.1 | 8.2 | 64.3 | 42.9 | 3.4 | 2.0 |
| 0.0625 ug/mL-ADB42 | 3762 | 4 | 24.7 | 7.2 | 0.0 | 5.2 | 63.8 | 35.7 | 3.4 | 1.8 |
| 0.125 ug/mL-ADB42 | 3980 | 4 | 25.3 | 7.4 | 0.0 | 4.8 | 63.8 | 35.1 | 3.4 | 1.7 |
| 0.5 ug/mL-ADB42 | 3121 | 4 | 25.0 | 9.7 | 0.0 | 6.2 | 40.6 | 31.7 | 2.0 | 1.4 |
| 2 ug/mL-ADB42 | 4995 | 8 | 26.9 | 18.0 | 1.9 | 8.0 | 29.5 | 43.3 | 1.1 | 1.7 |
| 4 ug/mL-ADB42 | 1440 | 4 | 36.7 | 27.7 | 5.0 | 13.9 | 38.8 | 51.6 | 0.9 | 1.9 |
| 8 ug/mL-ADB42 | 1287 | 4 | 54.7 | 56.2 | 4.1 | 13.2 | 40.0 | 39.2 | 0.6 | 0.9 |
| 16 ug/mL-ADB42 | 910 | 4 | 17.5 | 34.8 | 31.5 | 32.6 | 46.6 | 36.4 | 0.6 | 1.0 |

**Fig. 1:**
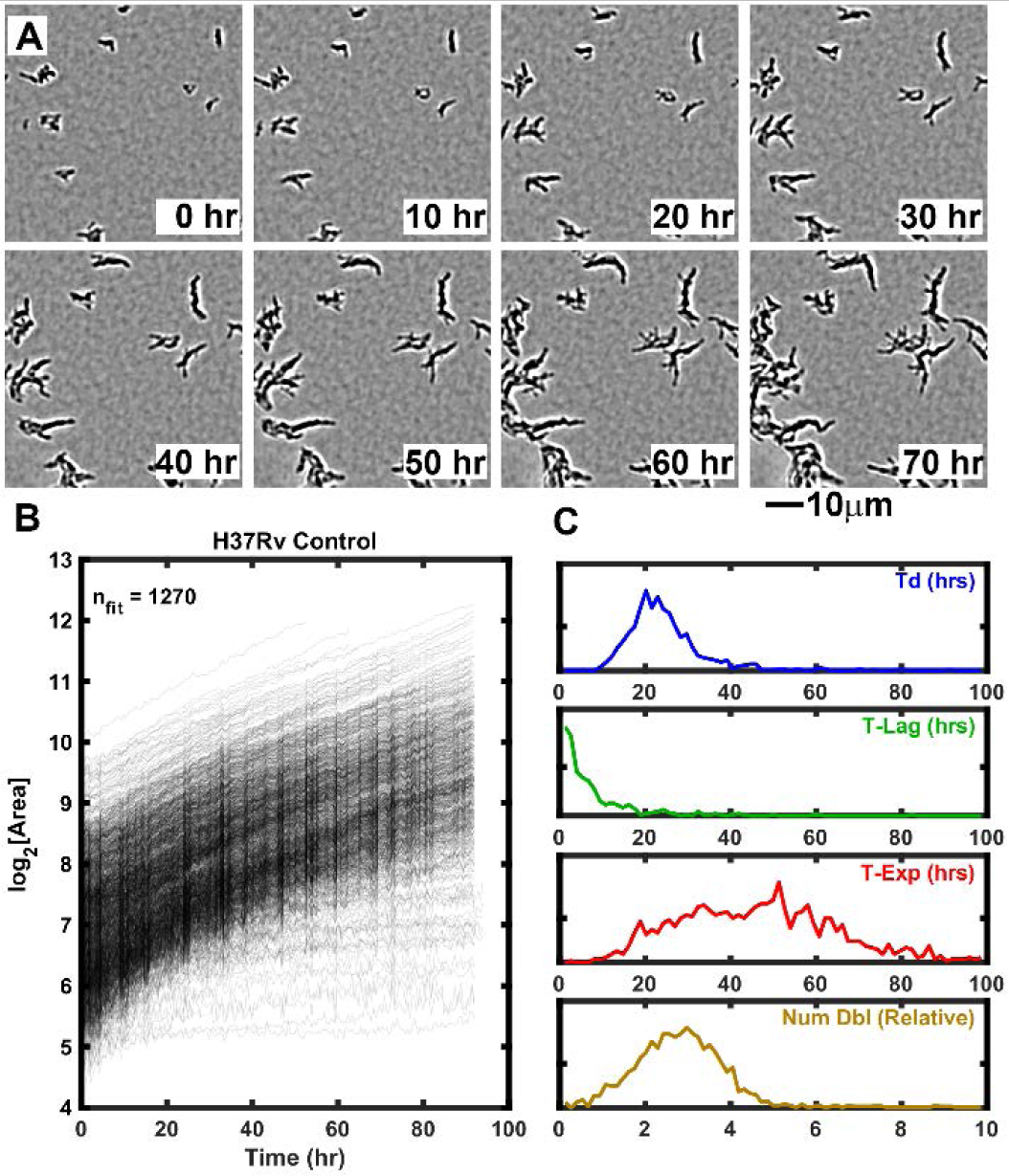
Growth of Mtb strain H37Rv into microcolonies over time observed with ODELAM. **A:** Selected time-lapse snapshots of MTB colonies growing on a solid medium. Mtb CFU are observed growing on a 60 μm x 60 μm region of 7H9-GO agar over 96 hrs time course. **B:** Growth curves for a total of 1276 colonies were recorded from a 1.2 mm x 0.9 mm region over 96 hours and plotted. **C:** Population histograms of the all extracted growth parameters (Doubling Time, Lag Time, Exponential Time and Number of doublings).

### Growth phenotype of Mtb strain H37Rv exposed to ofloxacin

One of the main advantages of ODELAM compared to standard bulk culture experiments is its ability to assess population responses on a single-CFU level. We therefore used ODELAM to reveal growth parameter values for individual CFUs in response to increasing drug pressures and compared these results to standard dose-response measurements (**Fig. 2**). We first focused on H37Rv. H37Rv is sensitive to ofloxacin with minimum inhibitory concentration (MIC) of 0.5 μg/ml as measured by normalized population growth in a standard dose-response curve (**Fig. 2B**). By ODELAM, we observed H37Rv to initially grow at all concentrations of OFX indicating that OFX’s toxicity does not alter growth kinetics over the first 24 hrs (**Fig. 2A**). This is consistent with OFX’s mechanism of action on the enzyme target, DNA gyrase, which leads to the accumulation of DNA damage (Manjunatha et al. 2002; Gore et al. 2006; Willmott et al. 1994). Above the MIC, toxicity led to cessation of growth, detected as an exit from exponential phase. This is reported as and a reduction in time spent in exponential phase (T-Exp) and in the number of doublings (Num Dbl) as shown by shifts of the population histograms (**Fig. 2C**). Narrowing of the population histograms reflects a more uniform population. The population distribution at the MIC remained relatively broad but was narrower than at lower drug concentrations or in controls. Notably, the doubling time distributions did not appreciably change; cells maintained similar doubling rates prior to growth cessation. Overall, the primary effect of OFX as detected by ODELAM was to arrest H37Rv growth after 2-3 doublings at the MIC and by the first doubling at high drug concentrations, while not appreciably affecting the rate at which H37Rv doubles prior to growth arrest (**Table I**).

**Fig. 2:**
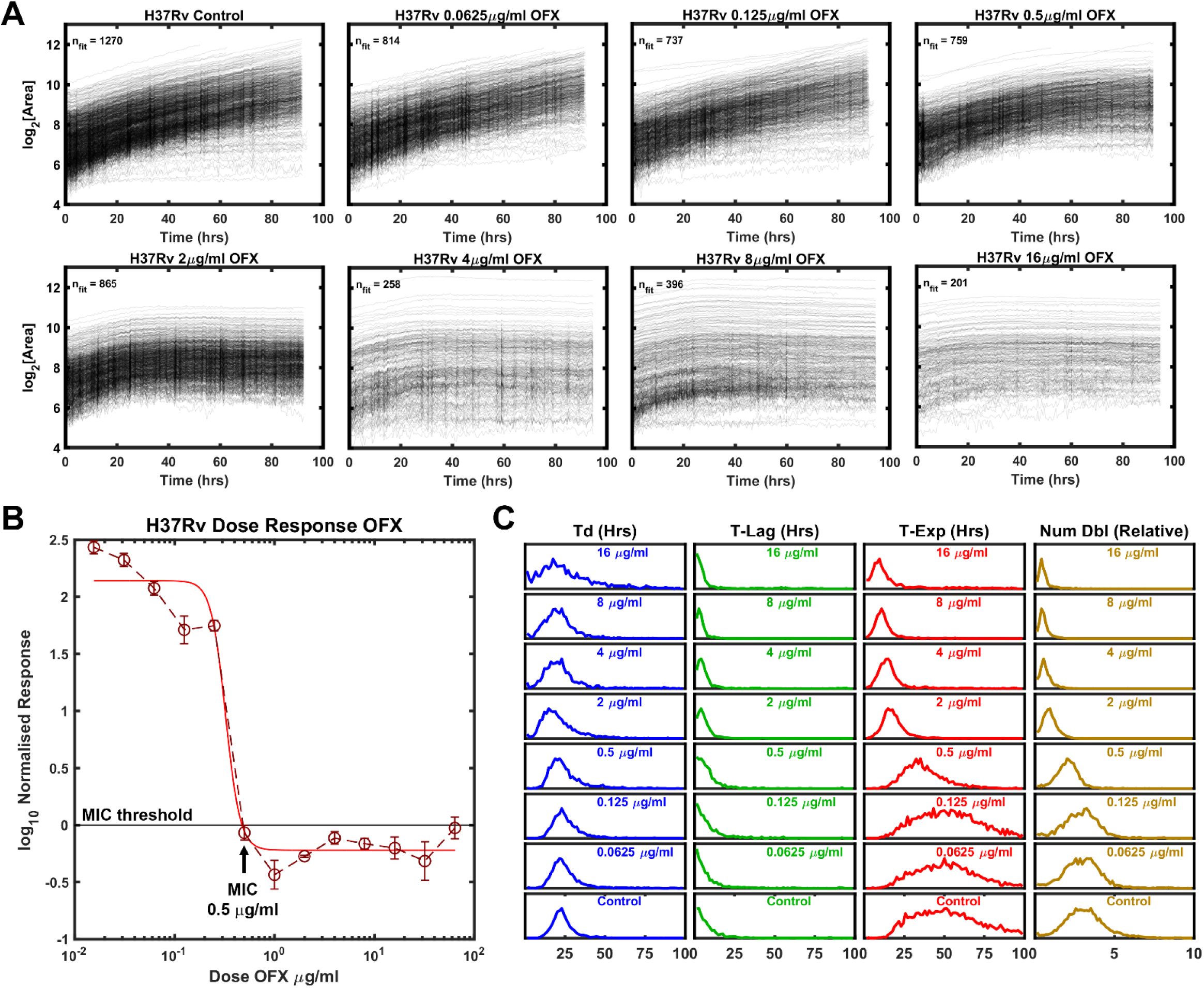
Growth of Mtb strain H37Rv during exposure to ofloxacin. A: Growth curves of H37Rv under increasing OFX drug concentrations measured by ODELAM. The flattening of the curves corresponds to the MIC of about 0.5 ug/mL OFX and corresponds to the batch culture dose response (**B**). **C:** Population histograms of the growth parameters showing that the time in exponential growth (**T-Exp**) is reduced as OFX concentration increases, which in turn leads to a reduced number of doublings.

### Growth phenotype of a resistant strain TRS10 exposed to ofloxacin

We observed a clinical isolate TRS10, which contains a missense mutation in the gene encoding gyrase A (D94G) that increases the isolate’s MIC to OFX (Eilertson et al. 2016; Hooper 2001). In the absence of drug pressure, the median doubling time of TRS10 was approximately 20% longer than that of H37Rv (**Table I**). The MIC of TRS10 to OFX was 16 μg/ml as measured by the normalized population growth in a standard dose-response curve (**Fig 3B**). By contrast to H37Rv, which ceased to grow at its MIC, TRS10 slowed its doubling time as OFX concentration increased up to, and including, its MIC (**Fig. 3A**). Further, TRS10’s doubling time slowed dramatically from ∼30 to ∼50 hrs as drug concentration shifted from 2 to 4 μg/ml **(Table I)**.

**Fig. 3:**
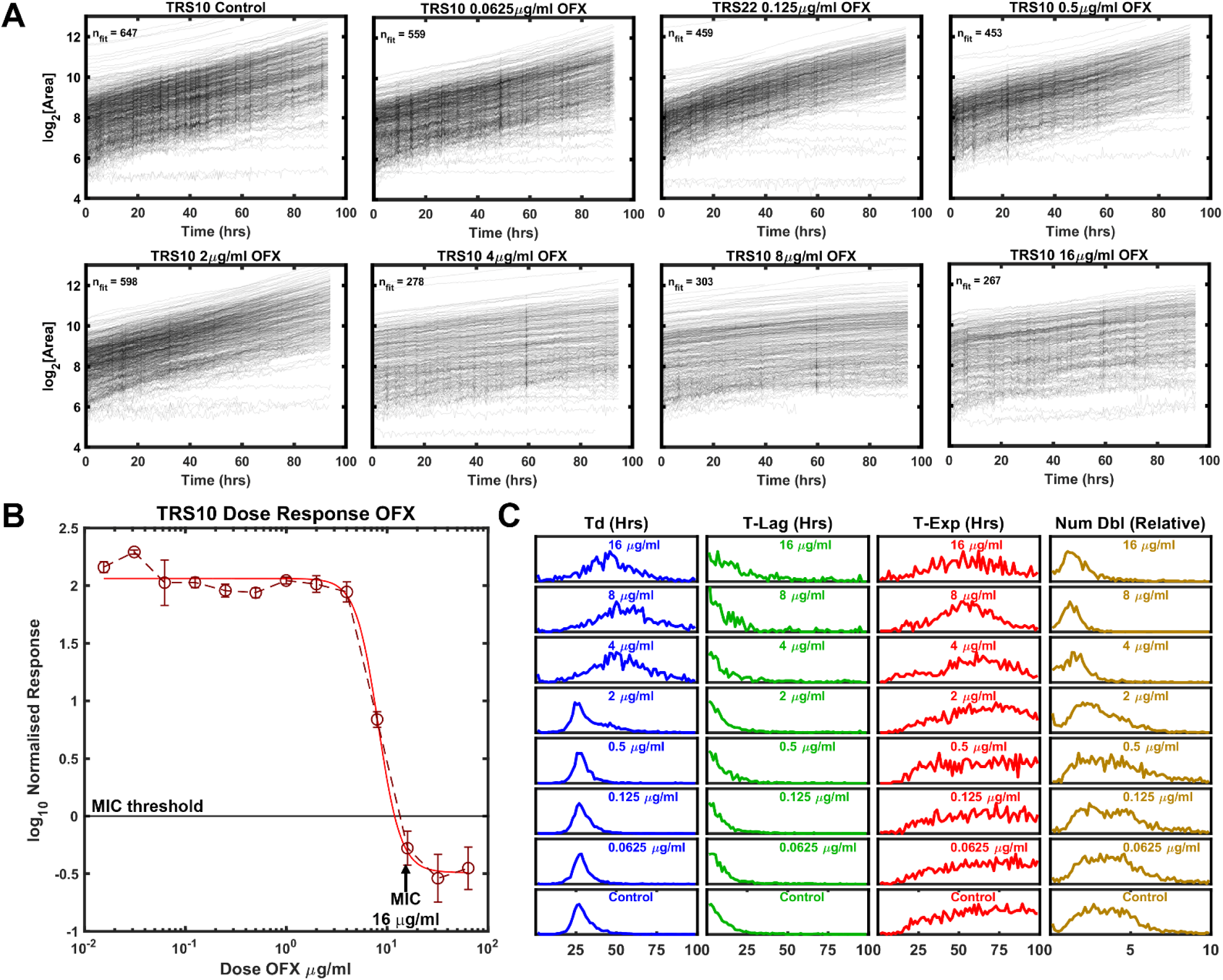
Growth of Mtb strain TRS10 during exposure to ofloxacin. **A:** Growth of clinical isolate TRS10 appears to slow under increasing OFX pressure. **B:** Batch culture dose response curve for TRS10, indicating an MIC of 16 ug/mL OFX. **C:** ODELAM histograms with increasing OFX concentration, showing that up to the MIC doubling time appears to increase while the time in exponential phase is less affected.

Total growth (Num Dbl) is influenced by all three growth parameters (Td, T-Lag and T-Exp). Therefore, to discern the contribution of these parameters to total growth we determined the population effect size, which quantifies the magnitude of effect for each parameter (Td, T-Lag, T-Exp, Num Dbl). The Kolmogorov-Smirnov statistic was used as a nonparametric score to evaluate the effect size and plotted against the concentration of OFX for each strain (**Fig. 4**) (Massey 1951). For H37Rv, the effect size for time in exponential phase tracked with the number of doublings as the concentration of OFX increased above 0.5 μg/ml while doubling time and lag time effect sizes did not (**Fig. 4A**). In contrast, for TRS10 the effect size for doubling time tracked with the number of doublings as the concentration of OFX increased (**Fig. 4B**). This OFX effect on TRS10’s doubling time as opposed to the cessation of growth observed for H37Rv (T-Exp) likely reflects the reduced binding of OFX to the mutant gyrase (Maruri et al. 2012; Willmott et al. 1994). Together these results show the ability of ODELAM and this analysis to reveal strain-dependent responses to antibiotics that would be missed by standard MIC assays. The examples of H37Rv and TRS10 responding to OFX (**Fig. 2** and **3**), as assayed by ODELAM, discerns these differing growth phenotypes and can inform on potential mechanisms of drug action.

**Fig. 4:**
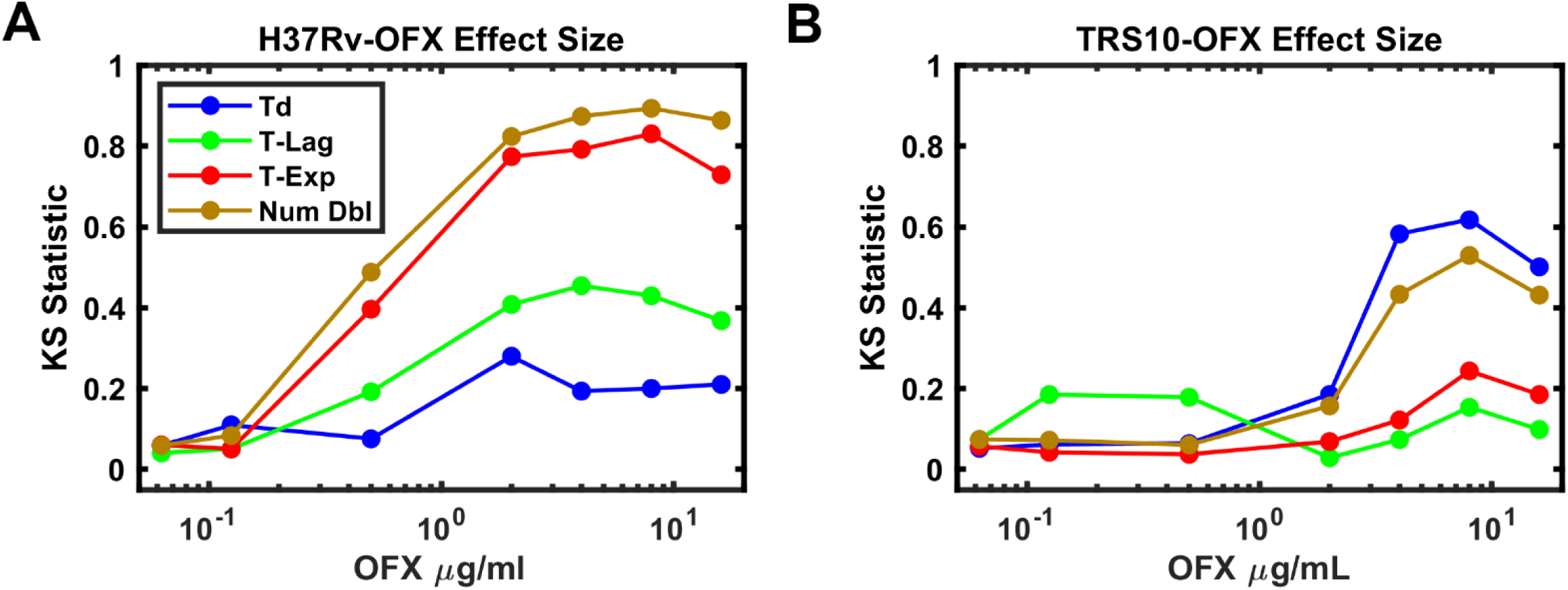
Effect Size of OFX on growth parameters. **A:** Effect size as measured by the Kolmogorov-Smirnov statistic vs OFX concentrations for H37Rv. Note, the number of doublings observed tracks with the time growing exponentially (T-Exp). **B:** Effect size as measured by the Kolmogorov-Smirnov statistic vs OFX concentrations for TRS10. Note, doubling time (Td) tracks with number of doublings (**B**).

### Detecting heteroresistance

Heteroresistance in clinical isolates is prevalent and confounds diagnosis and treatment of Mtb (Sebastian et al. 2016). Therefore, we tested the ability of ODELAM to rapidly detect heteroresistance. TRS10 was mixed with H37Rv to generate strain ratios ranging from 1:1 to 1:4166. Mixed cultures were assayed by standard dose-response batch culture MIC assay (**Fig. 5A**) and with ODELAM (**Fig. 5B and 5C**) at a concentration of 2 μg/ml OFX which discriminates OFX-sensitive and OFX-resistant CFU.

**Fig. 5:**
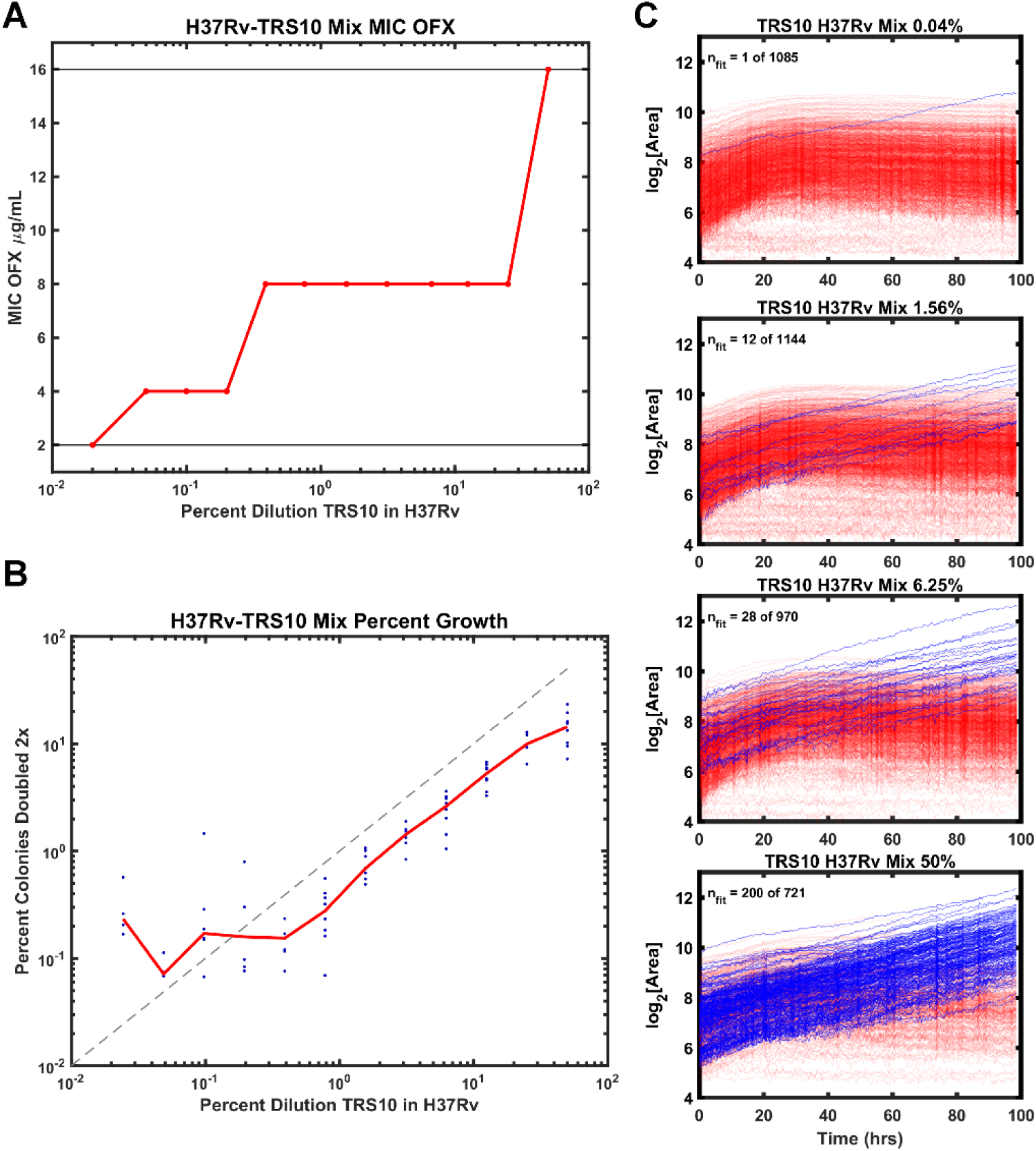
Comparison of ODELAM and MIC assay in detecting heteroresistance. **A:** Dilution of TRS10 into H37Rv and the corresponding MICs obtained from a bulk culture assay. **B:** Comparison of the percentage of TRS10 in H37Rv generated by dilution against the percentage CFU observed to grow more than 2x. The blue dots are replicate measurements from a given dilution and the red line is the mean of each replicate. **C:** ODELAM growth curves for TRS10/H37Rv mixes.

To evaluate the sensitivity of the MIC assay to heteroresistant cultures, the MICs of the dilution series were plotted (**Fig. 5A**). The observed MIC varied across the dilution series, increasing with the proportion of resistant cells. While the assay is sufficiently sensitive to detect shifts in the MIC at very low relative concentrations of resistant cells, it does not distinguish between a monoculture with a specific resistance phenotype and a mixed population with heteroresistance. Thus, the standard dose-response batch culture MIC assay yielded inaccurate results and was unable to reflect the MIC of the constituent population.

By contrast ODELAM directly observed OFX resistant CFUs in the mixed culture (**Fig. 5B,C**). Resistant and sensitive CFUs were distinguished by their number of doublings and their growth in exponential phase. Colonies that doubled more than two times and continued to grow in exponential phase for more than 30 hrs were considered OFX-resistant. In this ODELAM assay, a single OFX resistant CFU was detected from the 0.04% TRS10 mixture (**Fig. 5C**). Accordingly, there was an increase in the proportion of OFX-resistant CFUs detected as the percentage of TRS10 increased (**Fig 5B,C**). ODELAM consistently observed fewer resistant TRS10 CFU than expected by dilution. This may reflect differences in the ratio of OD600 to CFU for H37Rv and TRS10. The 50% mix detected 200 resistant CFU out a total of 721 CFU. Similarly, 28 of 970 CFU were detected in the 6.25% TRS10 mix, and 12 of 1144 were detected in the 1.56% TRS10 mix (**Fig. 5C**). A summary of the results from 12 dilution mixes and 8 replicates are plotted (**Fig. 5B**). The relationship between dilution and detection was linear from 50% to 0.2%. Below 0.2% the expected number of TRS10 CFU fall to around 1 to 5 in 1000 individuals and because each region of interest observes about 1000 CFUs, the measurements become dominated by chance.

### Detecting heteroresistance in clinical isolate ADB42

Clinical isolate ADB42 was previously observed to be OFX resistant with MIC ranging between 8 and 256 μg/ml of OFX (Eilertson et al. 2016; 2014). By whole genome sequence analysis, the canonical OFX resistance conferring gyrase-A D94G SNP was not present at a frequency sufficient to be associated with the observed OFX resistance; other mechanisms were attributed to the observed phenotype (Eilertson et al. 2016). In our hands, the standard dose-response assay yielded a MIC of 16 μg/ml (**Fig. 6A**). By ODELAM population measurements, the histograms revealed features indicative of phenotypic heterogeneity. Prominent tails were observed on the right-hand side of the exponential phase and number of doublings population distributions for OFX concentrations at and above 0.5 μg/ml (**Fig. 6B**), suggesting the presence of at least two populations. These populations could be segregated by a simple threshold of 2 doublings as detected by ODELAM on 2 μg/ml OFX (**Fig. 7**). The CFU growth curves corresponding to these populations showed either OFX sensitivity (**Fig. 7A**) or OFX resistance (**Fig. 7B**) similar to those observed for the experimentally mixed populations of H37Rv and TRS10 (**Fig. 5C**).

**Fig. 6:**
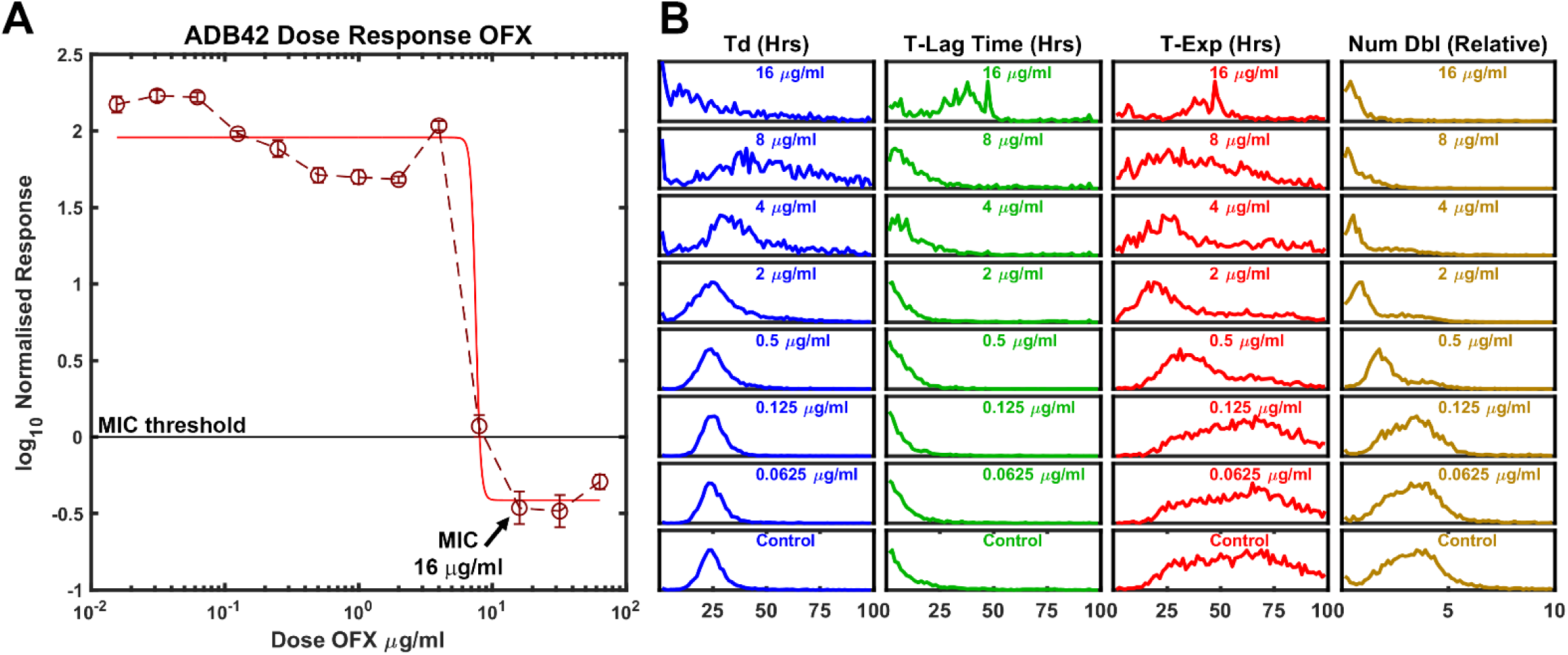
ODELAM analysis of Mtb strain ADB42 reveals heteroresistance to ofloxacin. **A:** Batch culture-based dose response curve of ADB42 showing an OFX MIC of about 16 µg/mL. **B:** At 2 µg/mL OFX tails in the doubling time, T-Exp, and number of doublings indicate that a second population may be present.

**Fig. 7:**
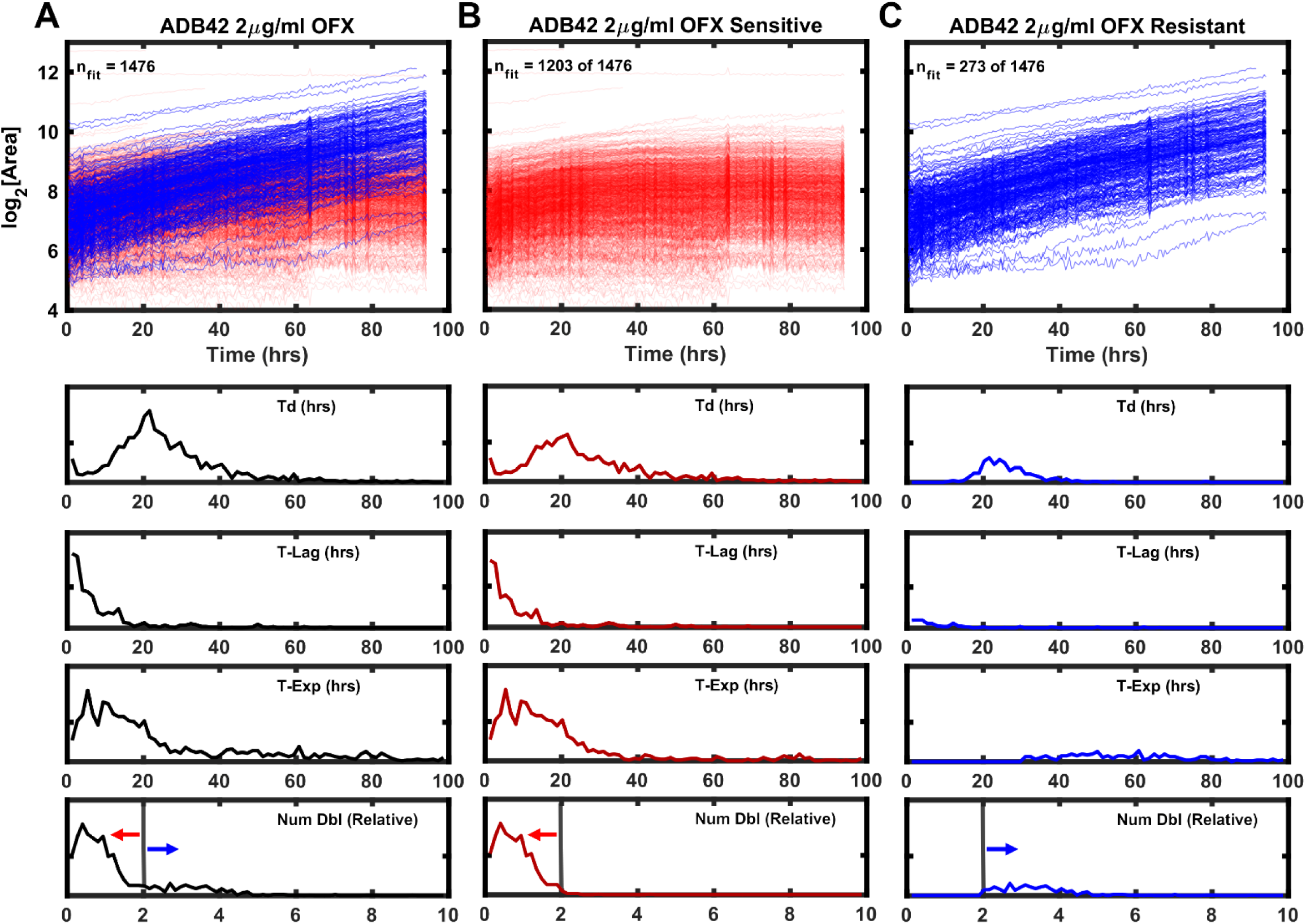
Growth kinetics generated by ODELAM for heteroresistant clinical isolate ADB42. **A:** Growth curves of ADB42 growing at 2 µg/mL OFX. **B:** Of the 1476 sensitive and resistant components of the heteroresistant culture, 1203 CFU may be segregated by selecting those that doubled fewer than 2x and, **C:** 273 CFU that doubled more than 2x as indicated by the gray line and the blue and red arrows.

ADB42 was grown in culture and subjected to transcriptome analysis by RNA-sequencing, which revealed the presence of a gyrase-A D94G SNP at a frequency of 15% (**Fig. 8A**). Across the experiments reported here, ODELAM measured roughly 24% of the ADB42 CFU were resistant to OFX, which is within a 95% confidence interval of random sampling modeled by a binomial distribution (**Table 2**). Ninety-six individual clones of ADB42 were isolated and their growth on 2 μg/ml OFX was measured in a spot assay (**Fig. 8B and 8C**). These clones were also measured with ODELAM to evaluate their growth kinetics and OFX resistance (**Fig. 9**). Thirty-three clones exhibited OFX resistance at 2 μg/ml OFX in both assays. In contrast to the parental clinical isolate, these individual clones did not display phenotypic heterogeneity by ODELAM (data not shown). In total, these data indicate that clinical isolate ADB42 contains two distinct populations one of which is resistant to ofloxacin and likely contains a D94G gyrase-A mutation that confers OFX resistance.

**Table 2:** Ratios of resistant cells as calculated by application of the binomial distribution and experimental observations

| **Total Number of Sample Reads (Number of Trials)** | **Number of D94G SNPs detected (Number of Success) $p = r$** | **Ratio of Resistant cells to sensitive cells (Probability of trial success)** | **95% Confidence Interval [Low]-[High]** |
| --- | --- | --- | --- |
| 62 | 9 | 0.09-0.24 | [2, 9] - [9, 21] |
| **Total Number of Colonies Picked (Number of Trials)** | **Number of Resistant Colonies (Number of Success) $p = r$** |  |  |
| 96 | 33 | 0.28-0.44 | [17,33] - [33,61] |
|  | **One or two cells contribute to colony $p = 1 - (1-r)^2$** | 0.14-0.26 | [17,33] - [33,61] |
| **Total Number of colonies tracked** | **Number that Fit Resistance Criterion** | **Ratio of Resistant cells to sensitive cells** |  |
| 963 | 237 | 0.2461 |  |
| 473 | 114 | 0.2410 |  |
| 1476 | 273 | 0.1850 |  |
| 1043 | 244 | 0.2339 |  |
| 359 | 101 | 0.2557 |  |
| 561 | 157 | 0.2799 |  |
| 573 | 146 | 0.2548 |  |
| 459 | 123 | 0.2680 |  |
|  | **Mean Resistance Ratio** | **0.2456** |  |

**Fig. 8:**
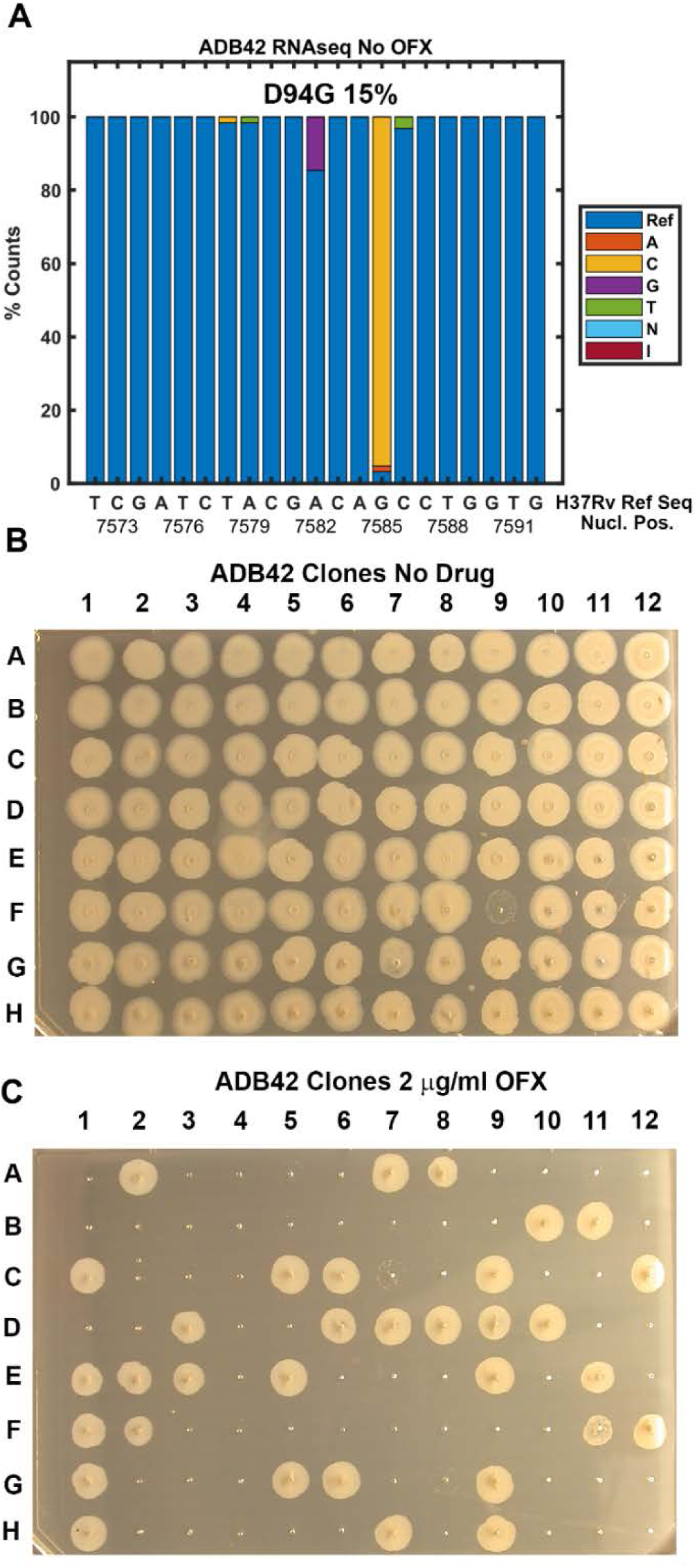
Assessment of ADB42 heteroresistance by spot assay. **A:** Prevalence of DNA gyrase II (Rv0006) D94G mutation before and after exposure to 32 μg/mL OFX for 24 hours as detected by DNA sequencing. **B&C:** Clones isolated from ADB42 grown on 7H10 media and 7H10 media with 2 μg/mL OFX for five days, before spotting on media lacking OFX **(B)** or containing OFC **(C)**. 33 of 96 clones were observed to grow in the presence of OFX, indicating resistance to OFX.

**Fig. 9:**
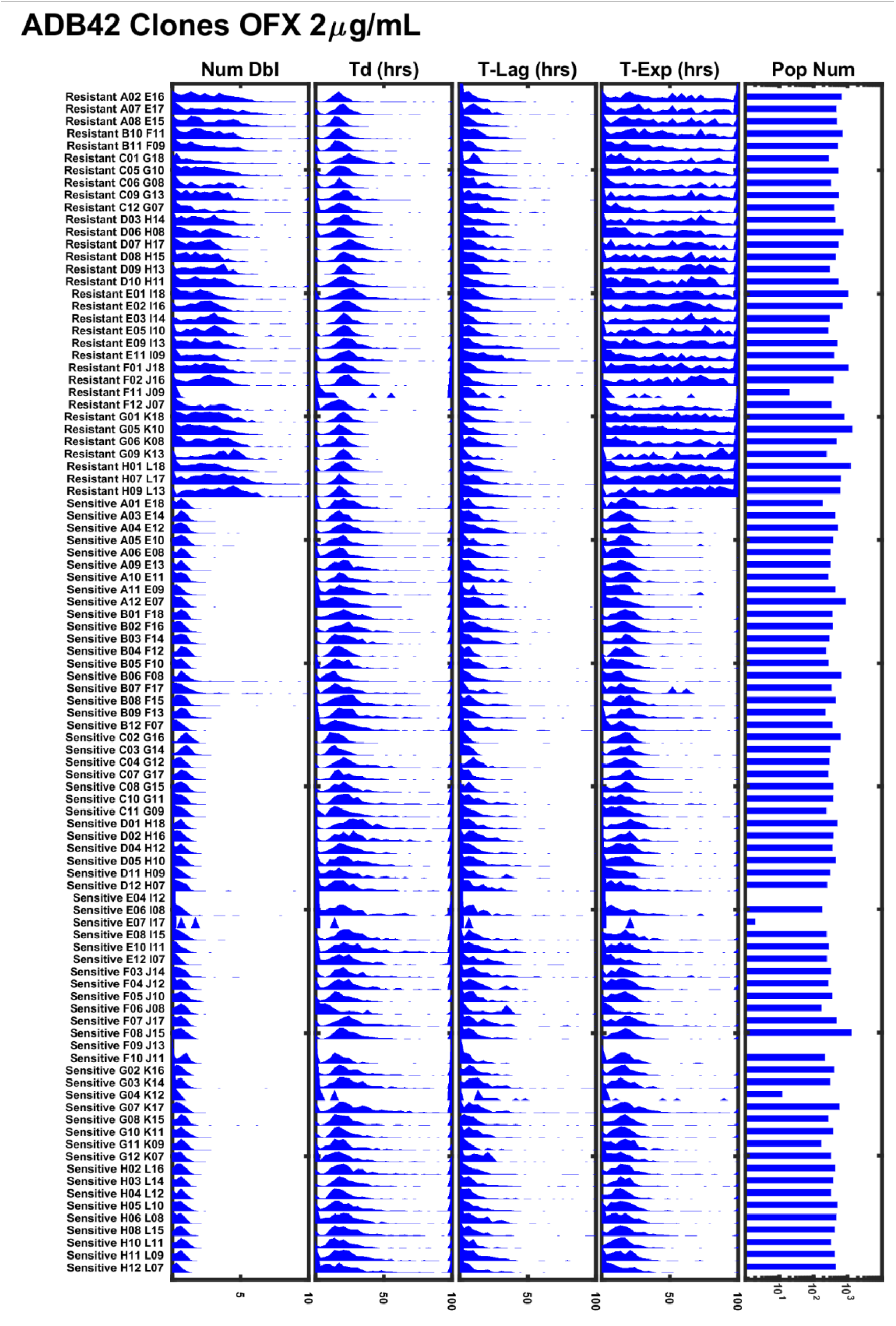
ODELAM experiment summary of 96 individual clones of ADB42 grown on 2 μg/mL OFX. Sensitive and resistant clones of ADB42 segregate according to their number of doublings (Num Dbl) and time in exponential phase (T-Exp). Sub-populations in the ADB42 clones were not observed.

In further support of these findings, we sequenced the quinolone-resistance-determining region (QRDR) of *gyrA* in fourteen of these ADB42 clones, 10 OFX-resistant and four OFX-sensitive. All 10 resistant isolates contained a SNP A7582G in *gyrA*, which corresponds to the canonical D94G mutations that confers OFX resistance (Willmott 1994). The four sensitive clones did not contain this SNP (**Table 3**). Additional gyrase mutations were present in the QRDR of *gyrA* but these mutations were not associated with OFX sensitivity or resistance (**Table 3**).

**Table 3:** Sequence data *gyrA* SNPs in OFX sensitive and resistant clones.

|  |  | QRDR | QRDR |  |
| --- | --- | --- | --- | --- |
| Position in the gyrase protein | **21** | **94** | **95** | **247** |
| H37Rv | E | D | S | G |
| TRS10 | Q | **G** | T | S |
| ADB42 OFX Sensitive | Q | D | T | G |
| ADB42 OFX Resistant | Q | **G** | T | G |

### Observation of OFX induced cell lysis

ODELAM directly observes cells and therefore informs on additional cellular phenotypes. After Mtb growth arrest on OFX, some cells appear to lyse as detected by a change in contrast (**Fig. 10A**), which we interpret as a consequence of cytotoxicity (**Fig. 10A**). This phenotype is observed as a sharp reduction in a colony’s size in its growth curve (**Fig. 10B**). The percentage of colonies that have this phenotype increased with OFX concentration and became apparent at concentrations greater than 0.5 μg/ml OFX. The frequency of lysis scales with the sensitivity of the strain to OFX. Here H37Rv showed as much as 95% of the tracked colonies exhibiting cell lysis, where as ADB42 and TRS10 showed 65% and 30%, respectively (**Fig. 10C**). These observations by ODELAM highlight the complex transition from a cytostatic to a cytolytic phase of the drug-organism interaction.

**Fig. 10:**
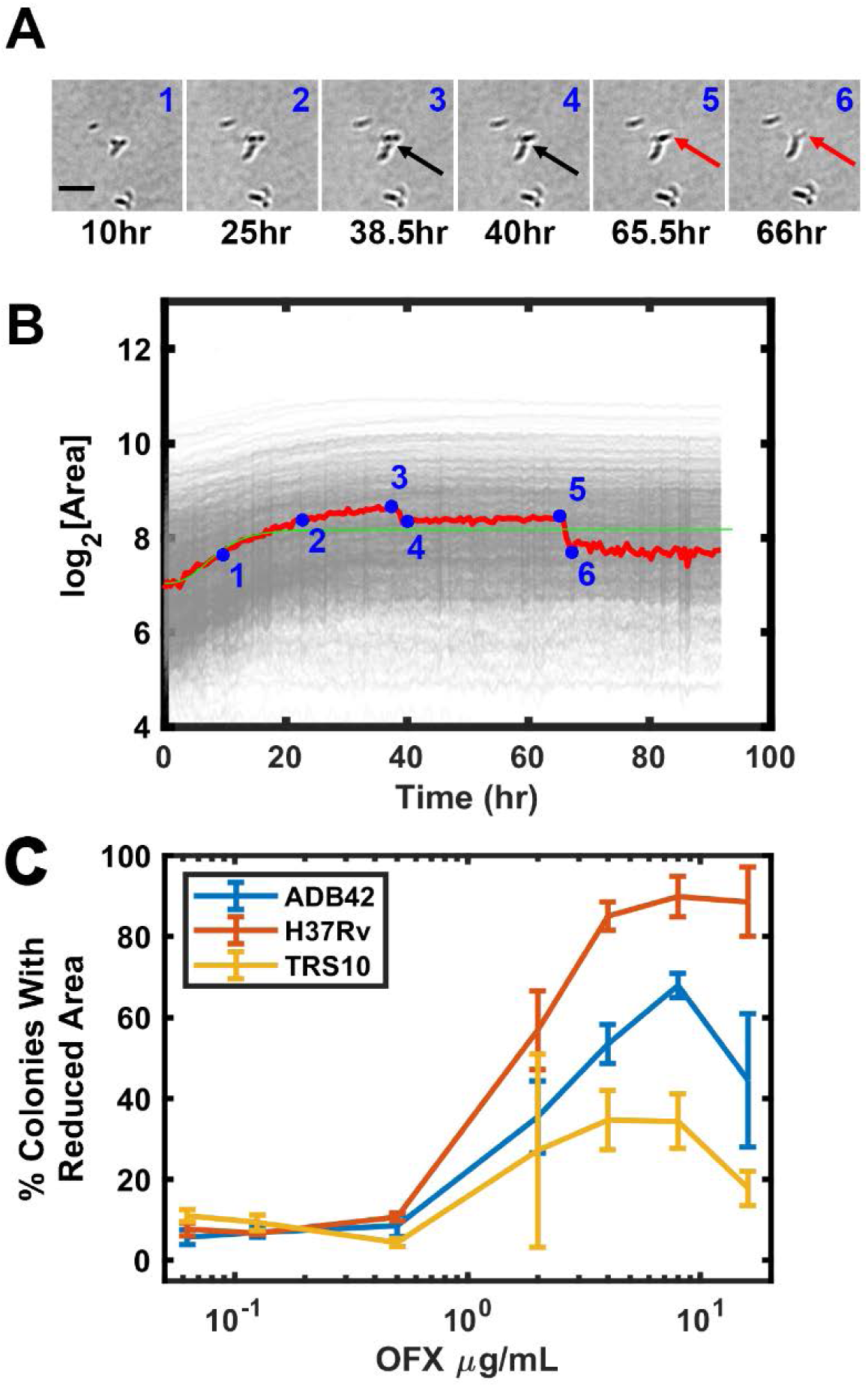
OFX induced lysis of the Mtb colonies. Some CFU were observed to lose area after exposure to OFX, which is attributable cell lysis indicated by arrows (**A**). Tracking of images in A and plotting CFU area over time detects lysis events as a reduction in CFU area (**B**). The percentage of colonies that lost area is plotted for each strain against drug concentration (**C**).

### Rate of Observation Time Convergence

Rapid detection of drug sensitivity improves in the successful treatment of tuberculosis. To establish the minimum time required by ODELAM to reliably identify anti-microbial resistance and sensitivity, we determined the length of time necessary to estimate a colony’s kinetic growth parameters. Each growth curve is fit using an algorithm optimized for ODELAM data. Ideal growth curves were generated in silico varying the kinetic parameters with random noise added. The Gompertz fit algorithm then estimated growth parameters for each noisy growth curve (**Fig 11A-C**). For each ideal growth curve, an increasing number of time points were included into the fit algorithm. As more time points from the ideal growth curve were added, the estimated parameters converged to stable values (**Fig 12A**). We evaluated how many time points needed to be included for the differences in sequential estimations to converge to a precision of 2.5% and 5% (**Fig. 12B-C**). In this way, we calculated the time required for reliably determining the kinetic parameters for a given growth curve, which is sensitive to each kinetic growth parameter. For the majority of cases, the ratio of time observed cannot be reliably shortened to less than the time taken for a CFU to exit exponential phase, which is dependent on drug sensitivity and noise in the measurement. Nonetheless ODELAM reliably detects OFX drug sensitivity of H37Rv within 30 hrs (**Fig. 2**).

**Fig. 11:**
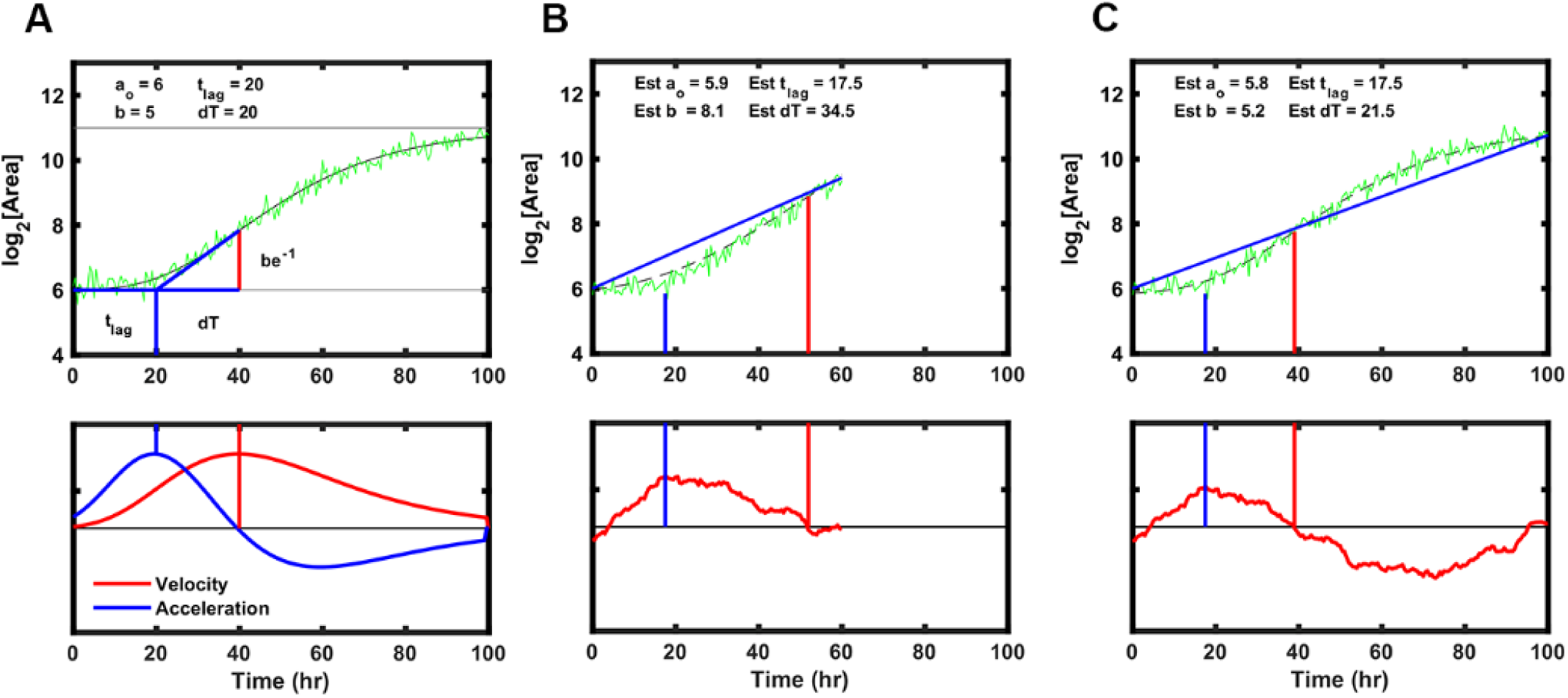
Strategy to fit Gompertz function parameters on a simulated growth curve simulated with noise added. **A:** Illustration of the parameters used to fit the Gompertz function as described in Materials and Methods. Note the value dT, the time between T-Lag and the maximum growth velocity, is used for estimating the growth curve but T-Exp is reported. T-Exp is the time between the maximum and minimum acceleration or twice the value dT. **B:** Initial estimates of each parameter were generated from a line connecting the first and last data points in a truncated simulated growth curve, which represents the time of experimental data acquisition. Bottom panel shows the differences between the line generated and the simulation. The initial estimate for T-Lag is determined by the maximum difference between the line generated and the simulation (blue vertical lines). T-Exp (or 2x dT) is determined where absolute difference between the line and the simulated data is minimized (red vertical line). **C:** Same as B with longer simulation time, showing initial estimates are similar.

**Fig. 12:**
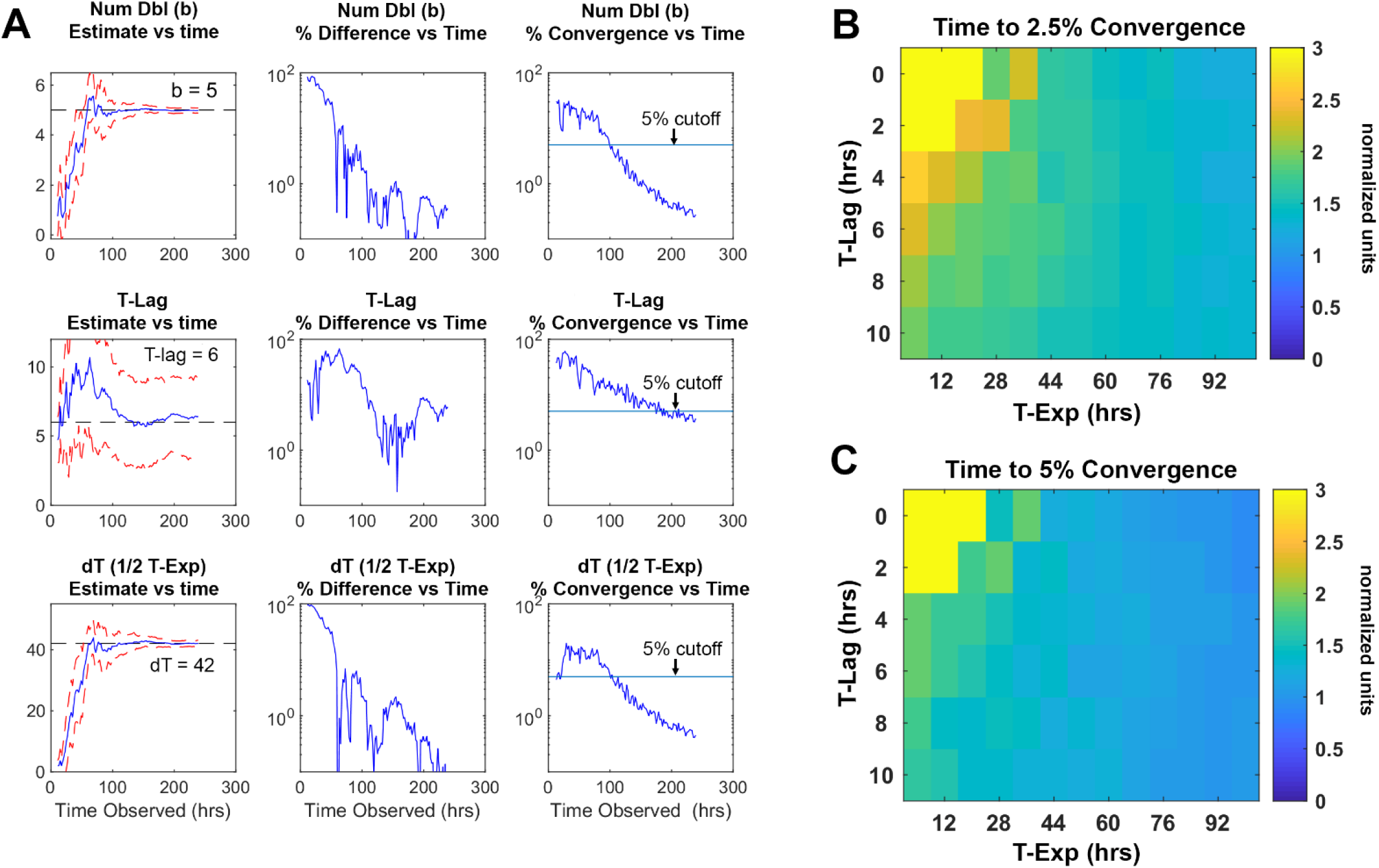
Time required for reliably determining the kinetic parameters in silico. **A**: Column 1, the difference between the true parameter, in this case Num Dbl = 5, T-Lag = 6, and dT = 42, and the estimated parameter are plotted as more of the growth curve is observed over time. As more timepoints of the growth curve are included or “observed” or included in the fit algorithm, the estimated parameters approach their true values. The red dashed lines are 1 standard deviation of 25 growth curves each with random noise added. Column 2, the percent difference between the true value and the estimate value are unstable and do not decrease consistently as more time points are included in the estimation. Column 3, percent convergence given by the difference between two consecutive time point estimations and divided by the true parameter value, and decreases steadily over time. The blue line represents a convergence threshold of 5% between consecutive estimations and this threshold is met when all parameters fall below the convergence threshold. **B-C:** Heat maps showing the ratio of time needed for the *T-Exp* parameter to converge to a given precision, *T*_*o*_ divided by the total time for the growth curve to begin to plateau *T-Lag* + *T-Exp*. The ratio is given explicitly as 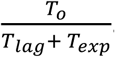. The heatmaps indicate for given lag times and time in exponential phase the parameters converge after the growth curve begins to plateau. This means that for a drug that causes growth cessation at 15 hours, a measurement of roughly 2x 15 or 30 hrs of observation is required for the *T - Exp* parameter to converge to a 5% precision.

## Discussion

As the rates of antimicrobial-resistant infections rise, the need for fast and unbiased assays to detect and characterize antimicrobial-resistant bacterial infections are required. ODELAM is a time-lapse, microscopy-based assay that detects Mtb drug sensitivity within 30 hrs, offers an unparalleled view into the population dynamics of Mtb growth kinetics and can directly observe phenotypic heterogeneity. While commercial multiplexed automated digital microscope-based techniques are available to rapidly screen for antibiotic resistance (Chantell 2015), here, using ODELAM we demonstrate the value of measuring growth kinetics on a single CFU level. With ODELAM we observed dramatically different effects of OFX on the growth parameters of drug sensitive and drug resistant clinical isolates of Mtb. These effects may offer insight into differing mechanisms of action for antimicrobials on Mtb or other organisms. ODELAM successfully discriminates between drug sensitive and drug resistant colonies in a mixed population. ODELAM also directly observes bacteriolysis and prolonged growth arrest, providing insight into the mechanisms of drug action. Based on this proof of principle, we suggest the investigation of additional clinical isolates, additional Mtb drug classes, and advances in this technology to improve throughput and feasibility in resource limited settings.

### OFX effects on growth kinetics

Growth kinetics are a powerful indicator of genetic fitness. ODELAM’s ability to monitor and evaluate growth kinetics of single CFUs from the point of plating through a time course of 100 hrs makes obtaining growth parameters and assessing drug sensitivity of multiple strains rapid and facile. Population growth kinetics under drug pressure illuminates the phenotypic differences between Mtb strains such as observed for the laboratory strain H37Rv, and an OFX resistant clinical isolate TRS10 (**Fig. 4**).

These two strains displayed very different phenotypic responses to OFX (**Figs. 2, 3 and 4**). H37Rv arrested growth at its MIC of 0.5 µg/ml as detected by departure from the exponential phase of growth. By contrast TRS10 grew continuously over a 96 hrs time period at its MIC OFX concentration of 16 μg/ml. Before growth arrest at its MIC, the doubling rate of H37Rv slowed slightly, whereas the doubling rate for TRS10 slowed considerably and at a population level exhibited a broad spread of doubling times. This suggests that the mechanisms of action for OFX toxicity are different between the two strains. As TRS10 contains the canonical D94G mutation in gyrase A conferring OFX resistance, we speculate that the growth phenotypes observed are due to the decreased affinity of OFX to the mutant DNA gyrase compared to wild-type, requiring a higher concentration of OFX to inhibit the enzyme. Thus, as the concentration of OFX is increased, off-target effects begin to manifest, which alter growth in a pleotropic manner resulting in the observed broadening of the doubling time of the population (**Fig. 3**). By contrast, inhibition of DNA gyrase in H37Rv likely results in the accumulation of replication and transcription intermediates, leading to the more uniform population distribution observed.

Growth kinetics of Mtb under fluoroquinolone antibiotic pressure have not been previously reported with the level of detail presented here. Fluoroquinolones such as OFX bind to gyrase and trap the enzyme-DNA complex in an intermediate state (Willmott et al. 1994). The trapped complex can stall RNA transcription and prevent progression of the replication fork (Willmott et al. 1994; Gore et al. 2006; Shea and Hiasa 1999; Manjunatha et al. 2002; Hiasa, Yousef, and Marians 1996). In the presence of OFX, double stranded breaks in the DNA template free the RNA transcription complex and accumulate over time contributing to toxicity and cell death (Willmott et al. 1994; Hiasa, Yousef, and Marians 1996). The time of exposure to OFX is thus directly linked to toxicity. Accordingly, even at OFX concentrations above the MIC (0.5 μg/ml), H37Rv continues to grow before they arrest synchronously after ∼20 hrs (**Fig. 2A**). We interpret this to mean that most cells remain metabolically active and synthesize protein until drug-induced damage accumulates and growth stops.

### ODELAM can directly observe heteroresistant subpopulations

Standard MIC assays are sensitive and robust methods for detecting antimicrobial resistant Mtb; however, these bulk assays give little information on the presence of sub-populations that differ in their sensitivity to drugs. ODELAM directly observes and quantifies such heteroresistance in a mixed culture, segregating populations according to their growth kinetics.

As ODELAM directly observes growth kinetics and transitions to stasis, it can differentiate drug sensitivity with precision not available in bulk assays. This was validated with an artificially mixed population and was used to detect heteroresistance in a clinical isolate, ADB42. In the mixed population ODELAM discriminates individual CFU with a sensitivity of roughly 1 in 1000. This level of precision is dependent on the number of CFU observed and improvements in microscope optics, computation, and media preparation could increase this sensitivity by orders of magnitude. Thus, this approach could observe the rare but critical events that lead to development of drug resistance including individual persistent cells that go on to acquire drug resistance through mutation (Cohen, Lobritz, and Collins 2013).

In the ADB42 isolate, OFX resistant cells were segregated by their growth kinetics in the presence of OFX. While DNA sequencing of this isolate failed to detect a resistant mutation, combining ODELAM with further DNA sequencing identified a D94G mutation in *gyrA* that we interpret conferred OFX resistance to a subpopulation in this clinical isolate. These observations are consistent with other measurements of strains with *gyrA* mutations (Eilertson et al. 2016).

### Observations of cytostatic and cytotoxic phases of drug-microbe interactions

ODELAM has the ability to directly observe cytotoxicity and the impact of drugs through different phases of growth. Remarkably, for concentrations of OFX up to and including 16 times the MIC, there was no measurable effect on H37Rv’s growth rate until growth ceased. By contrast OFX had a measurable effect on growth rate of TRS10 well below its MIC. We interpret these differences to reflect the differential accumulation of cytotoxic molecular intermediates due to differences in the ability of the drug to bind the different variants of gyrase. Nevertheless, ODELAM detected lysis of individual cells from each strain we analyzed in the presence of OFX at concentrations of 2 μg/ml and above (**Fig. 10**), providing a visualization of the onset of cytotoxicity. These measurements provide better assessments of drug action by resolving growth, cytostatic and cytotoxic states on a single cell level.

## Conclusions

ODELAM is a powerful time-lapse imaging technique for rapidly evaluating biological growth phenotypes in Mtb. The current methods integrate with common automated microscope equipment and are amenable to a Biosafety Level 3 environment. ODELAM directly observed heteroresistance in an Mtb clinical isolate and rapidly detected drug sensitivity in as little as 30 hrs. ODELAM is not restricted to Mtb and can be adapted to any colony forming microorganism. Insights from such studies promise an expanded view into the understanding of different mechanisms of antimicrobial resistance and drug action in Mtb and other important human pathogens.

## Materials and Methods

### Mtb Strains and Culture Methods

*Mycobacterium tuberculosis* was cultured as follows. Clinical isolates (Eilertson et al. 2016) were thawed and grown at 37°C to an optical density OD_600_ of 1.5 in 10 ml Middlebrook 7H9 media supplemented with glycerol, OADC supplement and 0.05 vol% Tween-80 (7H9-GOT) in 50 ml conical screw cap Falcon tubes. The samples were then stored at an OD_600_ of 1 in 0.5 ml aliquots of 15% (v/v) glycerol and frozen at −80°C until needed. For each experiment, a 0.5 ml aliquot was thawed and 9.5 ml of 7H9-GOT added. The culture was grown for 2-3 days until the OD_600_ was 0.3-0.5. The sample was diluted to 0.05-0.10 OD_600_ and grown for an additional 2 days to allow the culture to become well established in exponential growth phase. Finally, the cultures were diluted to 0.045 OD_600_ for spotting with ODELAM. Three strains were investigated in this study: H37Rv (laboratory standard and OFX sensitive), TRS10 (OFX resistant clinical isolate) and ADB42 (OFX heteroresistant clinical isolate). Strain names do not reflect their patient sources.

### Determination of the Minimum Inhibitory Concentration (MIC)

*M. tuberculosis* cultures were grown in 7H9-GOT medium at 37°C to a 0.3-0.5 OD_600_. The cultures were then diluted to 0.05 OD_600_ in fresh 7H9-GOT. A single 96-well plate was used for each strain. A 2-fold serial dilution of ofloxacin (0.015, 0.031, 0.062, 0.125, 0.25, 0.5, 1, 2, 4, 8, 16, 32, 64 µg/ml) was generated. Strains growing without drug were used as a 100% control and additional 1:100 dilution of each strain (to final OD_600_ 0=0.0005) in the drug-free medium was used as a 1% control. Wells containing only medium were used to measure background fluorescence. The plates were incubated at 37 °C for 7 days. After incubation, viable Mtb cells were quantified using two methods, luminescence by applying the BacTiter Glo™ Microbial Cell Viability Assay reagent (Promega, Madison, WI, USA) and fluorescence by applying alamarBlue reagent (Bio-Rad Laboratories, Hercules, CA, USA). A volume of BacTiter-Glo™ reagent equal to the volume of cell culture medium present in each well (20 µl) was added to each well. After 20 min of incubation at room temperature, luminescence was measured using the Omega plate reader. AlamarBlue was added in an amount equal to 10% of culture volume was added to each well of the 96-well plate. Plates were incubated in 37 °C for 9 hrs and the fluorescence was measure using Omega plate reader, with an excitation wavelength at 544 nm and emission at 590 nm. MIC measurements were performed in three technical and two biological replicates. MIC value was defined as the lowest OFX concentration that inhibited growth compared to growth of the 1% control.

### Mtb Cloning and Sequencing

Clinical isolates ADB42 and TRS10 were grown as previously described. After the culture reached an OD_600_ 0.5 the culture was diluted 10^5^, 10^6^, and 10^7^ CFU/ml and plated on 7H9 media in petri dish. Single colonies were then picked with a pipette tip and subcultured in a 100 μL of 7H9-GOT media in a 96 well plate. After growth to an OD_600_ of about 0.2 per well, cultures were diluted 10× and cultured at 37 °C for an additional 3 days. Then OFX resistance was evaluated by spotting 2 µl of clonal cultures were spotted onto 7H9 media with 0 and 2 μg/ml OFX. After two weeks of growth on agar plates were evaluated and photographed to determine resistant and sensitive clones. Additionally, the 96 clones were evaluated with ODELAM by spotting onto 7H9-GO with 2 μg/ml OFX and growth recorded for 96 hrs.

The quinolone resistance-determining region (QRDR) of gyrase A was amplified by PCR using the following method. ADB42, TRS10, and H37Rv were grown to an OD_600_ of 0.5 in 10 ml cultures. The cultures were spun and resuspended in 0.53 ml of TE buffer. Cells were mechanically lysed three times for 30 sec with 0.1 mm silica beads (Lysing Matrix B, MP Biomedicals). The samples were then spun, supernatant was transferred into new tubes, and heated for 30 min at 105 °C. DNA was purified using MagJet Genomic DNA kit (Thermo Fisher Scientific) according to the manufacturer’s instructions. DNA fragments for sequencing were amplified by PCR using a mix of 1 ng of genomic DNA, 5 pmol of each primer (gyrA_1Fw and gyrA_1Rv), 200 μM dNTPs, 1× Prime Star buffer and Prime Star polymerase. The primers utilized are given (**Table 4**). PCR products were purified using a NucleoSpin Gel and PCR clean up kit (Macherey Nagel). Sanger sequencing of PCR products were performed by GENEWIZ (Seattle, WA, USA).

**Table 4:**
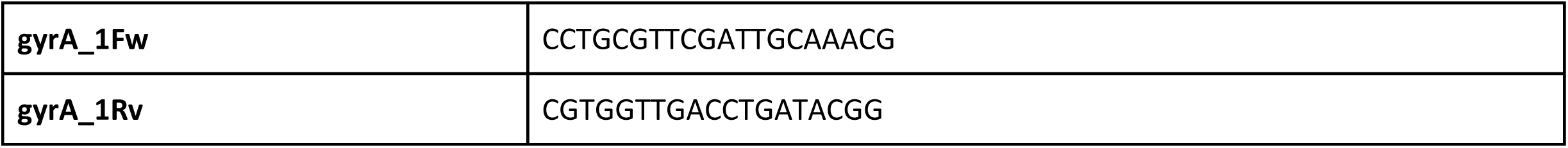
Quinolone Resistance Determining Region (QRDR) Primer Sequence:

### RNA isolation and Mtb transcriptome sequencing and analysis

RNA was isolated from these cultures as described previously (Sherman et al. 2001). Briefly, cell pellets in Trizol were transferred to a tube containing Lysing Matrix B (MP Biomedicals) and vigorously shaken at maximum speed for 30 sec in a FastPrep 120 homogenizer (QBiogene) three times, with cooling on ice between shakes. This mixture was centrifuged at maximum speed for 1 min and the supernatant was transferred to a tube containing 300 μL chloroform and Heavy Phase Lock Gel (Eppendorf), inverted for 2 min and centrifuged at maximum speed for 5 min. RNA in the aqueous phase was then precipitated with 300 μL isopropanol and 300 μL high salt solution (0.8 M Na citrate, 1.2 M NaCl). RNA was purified using a RNeasy kit following the manufacturer’s recommendations (Qiagen) with one on-column DNase treatment (Qiagen). Total RNA yield was quantified using a Nanodrop (Thermo Scientific).

To enrich the mRNA, ribosomal RNA was depleted from samples using the RiboZero rRNA removal (bacteria) magnetic kit (Illumina Inc, San Diego, CA). The products of this reaction were prepared for Illumina sequencing using the NEBNext Ultra RNA Library Prep Kit for Illumina (New England Biolabs, Ipswich, MA) according to manufacturer’s instructions, and using the AMPure XP reagent (Agencourt Bioscience Corporation, Beverly, MA) for size selection and cleanup of adaptor-ligated DNA. NEBNext Multiplex Oligos for Illumina (Dual Index Primers Set 1) were used to barcode the DNA libraries associated with each replicate and enable multiplexing of 96 libraries per sequencing run. The prepared libraries were quantified using the Kapa qPCR quantification kit, and were sequenced at the University of Washington Northwest Genomics Center with the Illumina NextSeq 500 High Output v2 Kit (Illumina Inc, San Diego, CA). The sequencing generated an average of 75 million base-pair paired-end raw read counts per library.

Raw FASTQ read data were processed using in-house R package DuffyNGS, as originally described (Vignali et al. 2011). Briefly, raw reads pass through a 3 stage alignment pipeline: 1) a pre-alignment stage to filter out unwanted transcripts, such as ribosomal RNA, mitochondrial RNA, albumin, globin, etc.; 2) a main genomic alignment stage against the genome(s) of interest; 3) a splice junction alignment stage against an index of standard and alternative exon splice junctions. All alignments performed with Bowtie2, using command line option ‘--very-sensitive’ (Langmead and Salzberg 2012). BAM files from stages 2 & 3 are combined into read depth wiggle tracks that record both uniquely mapped and multiply mapped reads to each of the forward and reverse strands of the genome(s) at single nucleotide resolution. Multiply mapped reads are pro-rated over all highest quality aligned locations. Gene transcript abundance is then measured by summing total reads landing inside annotated gene boundaries or exon boundaries, expressed as both RPKM and raw read counts (Wold and Myers 2008). Two stringencies of gene abundance are provided, using all aligned reads and by using just uniquely aligned reads.

### Preparation of ODELAM agarose plates

A Ninjaflex 2 mm thick gasket was 3D printed onto a 50 mm × 75 mm × 1 mm glass slide (VWR) using a Makerbot 2× printer (Appendix I). Following printing, the slides were cleaned of particles and fibers using lab tape. An additional slide was cleaned with ethanol and placed over the gasket so the space between the slides defined by the gasket could be filled with molten agarose media. The assembled agar mold was held together with binder clips.

Agarose was prepared as described previously (Herricks et al. 2017). Bulk agarose was prepared by dissolving 2 g of agarose in 150 g of 18 MΩ H_2_O. The solution was microwaved in 15 sec intervals for about 3 min or until the agarose boiled and all agar particles were dissolved by visual inspection. The 3.1 g of molten agarose was then aliquoted into 15 ml falcon tubes and stored at 4 °C. On the morning of an experiment, 0.4 ml of 10x 7H9-G media, 0.2 ml of sterile 18 MΩ H_2_0 were added to five previously prepared 3 g aliquots of agarose tubes. The solution was boiled for 8 min in a covered boiling flask. After fully melting the agarose, the solution was placed into a 45 °C stirred bath to cool for about 2-10 min. Once cooled, 0.4 ml of Middlebrook OADC and 4 μl of 1000× concentration of drug or drug vehicle was added to make a total of 4 ml of 7H9 glycerol oleate (7H9-GO) agarose media. The media was then injected into the single or each of the 5 glass-gasket-glass chambers using a 20-gauge syringe needle. The agarose was allowed to set for about 30 min and one glass slide was removed to expose the agar surface. The agarose slide was then placed in a sterile tip box and stored at room temperature for about 2 hrs before spotting Mtb cultures.

### ODELAM Time-Lapse Microscopy

The agar plate was then assembled into the ODELAM growth chamber and 1 μL of 0.045 OD_600_ Mtb culture spotted on the agar plate. The chamber was assembled and electrical contacts placed on an Indium Tin-Oxide (ITO) coated cover-slide. The ITO slide was resistively heated with 10 V and 0.2 A of current to minimize condensation above the agar. The spotted CFUs were imaged using on a Nikon TiE microscope equipped with an In Vitro Scientific incubator maintaining a temperature of 37 °C. Images were recorded using a 20× 0.45 NA long working distance lens with the correction collar set to 1.3 mm. This collar setting was required to get a monomodal focus function as calculated using the Laplacian Variance Method (Pertuz, Puig, and Garcia 2013). A Photometrix CoolSnap ES^2^ camera recorded brightfield images. The microscope was controlled by MATLAB Graphical User Interface software using the MicroManager Core API (Edelstein et al. 2014). At each location where culture was spotted, the microscope would tile a 3 × 3 array with 20% overlap. Each spot would be imaged every 30 min for up to 120 hrs. The resulting images were then stitched, binarized with a threshold, and objects tracked using a MATLAB Image processing pipeline. Results were then quantified and plotted using MATLAB. ODELAM is optimized to track between approximately 50 to 1500 individual cells and CFUs for 96 hrs. The lower limit of about 50 CFU per field of view is required to have sufficient contrast to focus and stitch the images.

### Extraction of Growth parameters

Growth curves of log_2_ area over time were fit to a parameterized version of the Gompertz function:

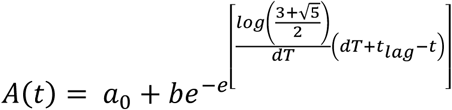

Where *a*_0_ is the initial area of the cells, *b* is the number of doublings the cells undergo at quiescence, *t*_*lag*_ is the lag time and *dT* is ½ the time the cells are growing exponentially T-Exp. Estimation of growth parameters was performed using the following strategy. The Gompertz function was written such that lag time *t*_*lag*_, the number of doublings *b*, and *dT* which is the time between the maximum velocity and *t*_*lag*_ both variables correspond to the maximum of the growth curves 1^st^ and 2^nd^ derivative respectively (**Fig: 11A**). This strategy allows the constraints *t*_*lag*_+ *dT* are less than or equal to the total time observed and that the value *b* is calculated from the observed area at the time *t*_*lag*_+ *dT*(**Fig: 11A**). Initial values for growth parameters were found using a either a geometric approximation of initial parameters or a course grid search. The geometric approximation was estimated by drawing a line between the first and last points of the curve. The residuals of that line approximate the sec derivative of the growth curve where the maximum value is approximately the lag time and where the residuals cross zero corresponds to the growth curves maximum velocity (**Fig: 11B and 11C**). For ease of analysis the growth curves were smoothed with a moving average window that was 5 time points wide. Course grid optimization routine would be used if the geometric approximation failed. The initial parameter estimates were then optimized using the MATLAB function fmincon. Doubling time was derived from the three parameters *b* and *dT*. All data was plotted using MATLAB software. A threshold criterion for including colonies for data analysis was that the colony was tracked for greater than 30 hrs, and those that showed a log_2_ growth greater than 0.1 were plotted.

### Statistical analysis of ODELAM data

Effect size was measured by pooling replicate measurements into a single population. The resulting population distributions were compared using the 2-sample Kolmogorov-Smimov test against a no-drug control pool. The KS test statistic for each comparison was then plotted to show the trend in OFX concentration vs. effect size. The binomial distribution was used to estimate the percentage of resistant cells in clinical isolate ADB42. For clarity the binomial probability distribution function is given by:

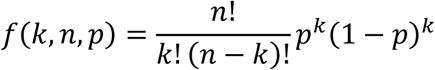

where *n* is the number of resistant colonies or SNP reads, *k* is the total number of colonies picked or the total number of reads at SNP locus and *p* is the fraction of resistant CFU in the culture. The binomial cumulative distribution function was calculated using the MATLAB function cdf. In this way we could model the number of colonies that appear resistant or the number of whole genome sequencing reads that contain the D94G mutation in Rv0006 as a function of the percentage of cells in the culture that are OFX resistant. The probability of success (*p*) is given by the percentage of OFX resistant cells which is estimated by the number of successful trials (given by number of SNP reads or the number of resistant colonies) is bounded by the 95% confidence interval that still contains the actual number of WGS reads or resistant colonies (i.e. number of successes) observed.

## Appendix I

### Description of Chamber and 3D printed Gasket

STL file for gasket description of steps needed to offset G-code to print on elevated slide.

Mechanical drawings and STL files of chamber.

### Description and figure of ODELAM growth chamber

Files for design of the chamber are available

### Abbreviations

CFU: colony forming units
MIC: Minimum Inhibitory Concentration
Mtb: *Mycobacterium tuberculosis*
Td: doubling time
T-Lag: Lag time
T-Exp: Exponential growth time
Num Dbl: Number of doublings
OFX: ofloxacin
ODELAM: One-cell Doubling Evaluation of Living Arrays of Mycobacterium
KS: Kolmogorov-Smirnov
GOT: glycerol oleate tween

